# The La Crosse virus Gc head domain is a major determinant of virus dissemination and pathogenesis

**DOI:** 10.1101/2025.05.12.653523

**Authors:** Ariana Dedvukaj, Nicole C. Rondeau, Matthew C. Lutchko, Kenneth A. Stapleford

**Affiliations:** Department of Microbiology, New York University Grossman School of Medicine. New York, NY 10016

## Abstract

How orthobunyaviruses establish infections and disseminate to cause disease is not well understood. In a previous study using the *in vivo* evolution of La Crosse virus (LACV), we discovered a cluster of mutations localizing to the LACV Gc head domain. However, we do not understand how the Gc head domain contributes to infection. We generated each of the aforementioned mutations and addressed the role of the Gc head domain in viral replication and infectivity in human neurons and myoblasts. We found that specific head domain residues could attenuate replication and infectivity in both cell lines, indicating an important role for the head domain during infection *in vitro.* Focusing on the Gc N609D variant which was attenuated *in vitro*, we infected three-week-old WT C57BL/6J mice via the footpad with WT LACV or the Gc N609D variant and found that the Gc N609D virus was completely attenuated. To address whether the variant was also attenuated in a highly susceptible mouse model, we infected *Ifnar1^-/-^* mice with WT LACV and Gc N609D and found that virulence in mice infected with Gc N609D was delayed with several mice surviving the infection. Subsequent studies looking at virus dissemination to the brain show that the Gc N609D virus has decreased neuroinvasive events, supporting a role for the head domain in virus dissemination. Together, these studies define a critical role of the Gc head domain in infectivity, dissemination, and pathogenesis. Studies are underway to further define how the orthobunyavirus Gc head domain contributes to infection and disease.

**Importance:** Orthobunyaviruses are emerging arboviruses capable of severe disease and explosive outbreaks. However, our understanding of how orthobunyaviruses establish infections or cause disease is not completely understood. The orthobunyavirus Gc glycoprotein contains a variable amino-terminal head domain that forms the tip of the virion trimeric spike, yet it is unclear how the head domain contributes to infection or pathogenesis. In this study, we use La Crosse virus (LACV) and a panel of Gc head domain variants to address the role of the head domain in LACV biology. We found that critical head domain regions are important for virus infectivity and pathogenesis in mice, highlighting an important role for the Gc head domain in orthobunyavirus infection and disease.

## Introduction

The orthobunyavirus genus (*Peribunyaviridae*) includes a long list of significant human pathogens (1, 2). These arthropod-borne viruses (arboviruses) are transmitted to humans primarily by mosquitoes, midges, and ticks. Recent outbreaks of Oropouche virus (OROV) in Cuba (3) and South America (4–7) along with the prevalence of La Crosse virus (LACV) in the United States emphasize the clinical relevance of orthobunyaviruses. LACV is mainly found in the East North Central and Appalachian regions (8). A member of the California serogroup of orthobunyaviruses, it is related to other neuroinvasive human viruses found worldwide, including Jamestown Canyon virus, Inkoo virus and Tahyna virus (9). Although the majority of cases are asymptomatic and therefore underreported, LACV is the leading cause of pediatric arboviral encephalitis in the United States. Neuroinvasive disease can result in fatality or lifelong neurological sequalae, such as recurring seizures and cognitive deficits (10) (11) (12). However, there are currently no antiviral therapies or vaccines targeting orthobunyaviruses, highlighting the need to study orthobunyavirus biology in molecular detail.

Our limited antiviral therapies are in part due to our incomplete understanding of the molecular mechanisms orthobunyaviruses use to establish infections. The orthobunyavirus negative-sense RNA genome consists of an S, M, and L segment. The S segment encodes the nucleoprotein and interferon antagonist, NSs. The M segment encodes the Gn and Gc glycoproteins along with non-structural protein NSm (1, 13), and the L segment encodes the RNA-dependent RNA polymerase. The M segment proteins are important for virion assembly (13, 14), entry (15–17), and cell-to-cell spread (18), yet we know little of how discrete domains within these proteins contribute to virus infection. Specifically, the orthobunyavirus Gc protein is a class II fusion glycoprotein similar to those of other bunyaviruses as well as alpha- and flaviviruses (19–22). Orthobunyavirus Gc contains a unique amino-terminal variable head domain that forms the tip of the glycoprotein spike (23, 24). Previous work with Bunyamwera virus and OROV has shown that the head domain can be deleted with little impact to virus replication *in vitro* and that fusion and immunogenicity is enhanced, suggesting that the head domain plays an important yet unknown role in virus biology (25, 26). However, even considering observations, we do not understand specifically how the Gc head domain contributes to infection or pathogenesis.

In this study, we took advantage of *in vivo* virus evolution which identified the La Crosse virus (LACV) Gc head domain as a domain for virus adaptation (15). Using these studies, we generated a panel of LACV Gc head domain variants and addressed the role of these residues and the Gc head domain *in vitro* and *in vivo*. We found that specific Gc head domain residues could influence replication and infectivity both *in vitro* and *in vivo*. Focusing on the most attenuated Gc variant N609D, we found that this variant could completely attenuate virulence of a pathogenic WT LACV strain in wild-type mice. Moreover, we found that Gc N609D was critical for dissemination and virulence in *Ifnar1^-/-^* mice. Finally, using an evolutionary approach we found that the LACV Gc head domain has been evolving across lineages and that there are conserved elements maintained within the head domain of related orthobunyaviruses. Together, these studies provide a critical role for the orthobunyavirus Gc head domain in virus infectivity, dissemination and virulence and allow us to speculate on how changes in the orthobunyavirus head domain may impact virus biology.

## Results

### The LACV Gc head domain influences virus growth *in vitro*

The orthobunyavirus Gc glycoprotein amino-terminal variable head domain sits atop of the glycoprotein spike (22–24). However, it is unclear how the head domain contributes to LACV infection *in vitro* or *in vivo.* In a previous study, we used *in vivo* evolution of La Crosse virus (LACV) in *Aedes (Ae.)* mosquitoes and C57BL/6J mice to identify potential residues that may be important for virus infection (15). We identified eight head domain variants which we hypothesized were important for LACV infection (**Fig. 1A and B**). Each of these residues is located at the head domain interface between Gc molecules (15, 24) (**Fig. 1A**). Importantly, when we aligned the M segment protein regions deposited in NCBI, we found several of these variants (W618R, D619G, and E623G) have been found in nature, highlighting their potential role in LACV infection (**Fig. 1B**). Given the location of these residues, we hypothesized that the head domain is critical for virus infection. To test this hypothesis, we generated each head domain variant in the LACV Lineage I Mosq/NC/1978 infectious clone M segment (27). All variants were genetically stable after passaging virus once in Vero cells to generate working stocks. To begin, we looked at plaque size of each variant on Vero cells. We observed that the variants Gc N609D and W618L led to small plaques while variants W618R, D619G, A621V, and E623A led to larger plaques compared to wild-type (WT) LACV (**Fig. 1C**).

**Figure 1:**
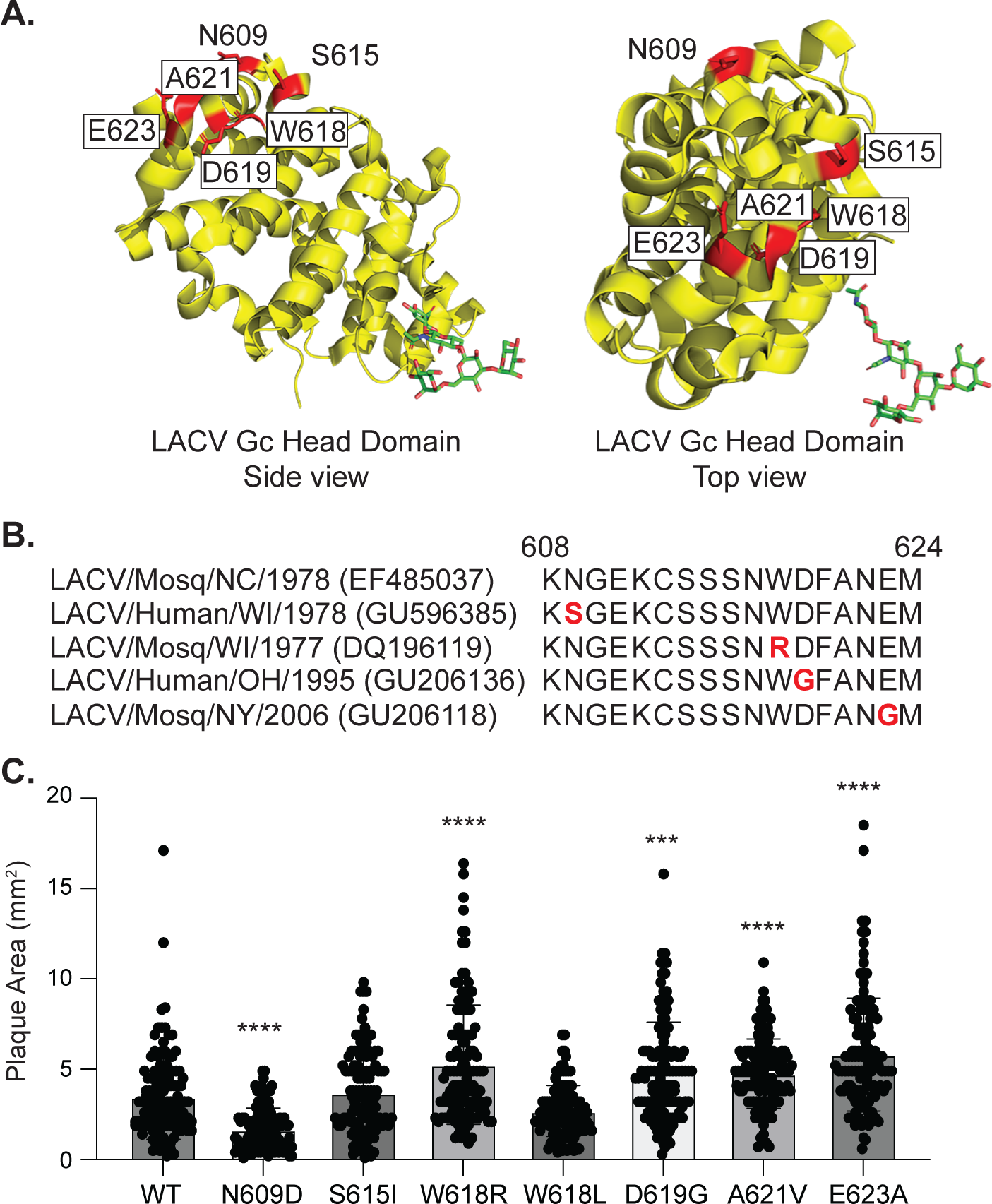
La Crosse virus Gc head domain variants influence plaque size. **A.** Crystal structure of the LACV Gc head domain (PBD: 6H3W) side view and top view with variants shown in red. **B.** LACV sequence alignments of the Gc head domain variant amino acid region 608 to 624. Accession numbers are located to the right of each name. Natural variants are shown in red. **C.** Plaque size of wild-type LACV and each Gc head domain variant on Vero cells 72 hours post-infection. Results represent the mean and standard deviation (SD) of two independent infections. N=120 plaques for each virus. Kruskal-Wallis test. *** p < 0.001, **** p < 0.0001.

To study the role of each residue in LACV replication *in vitro*, we first performed multi-step growth curves in Vero cells (**Fig. 2**). We used Vero cells specifically because they are deficient in interferon signaling, allowing us to focus on the role of each Gc residue without confounding influence from the antiviral response. In Vero cells, the replication of the Gc variants separated into two groups. We found that several of the variants including Gc N609D, S615I, and W618L led to attenuated growth; while D619G, A621V and E623A led to enhanced growth over WT LACV (**Fig. 2A**), suggesting that the Gc head domain can both enhance and attenuate virus growth *in vitro*. Given that several of the variants were selected for in *Ae. aegypti* mosquitoes, we also addressed viral growth in Aag2 *Ae. aegypti* cells (**Fig. 2B**). We observed that while several of the variants were attenuated early during infection, all of the variants caught up to WT LACV by 24 hours. When we sequenced each variant at 24 hours from the Aag2 cells, we found that all of the variants had reverted to the wild-type residue, suggesting that these variants may be restricted early during infection but a strong selective pressure forces them to revert to the WT residue. Together, given significant changes in vero cells and reversion in Aag2 cells, these studies show that the Gc head domain plays an important role in replication in mammalian cells.

**Figure 2:**
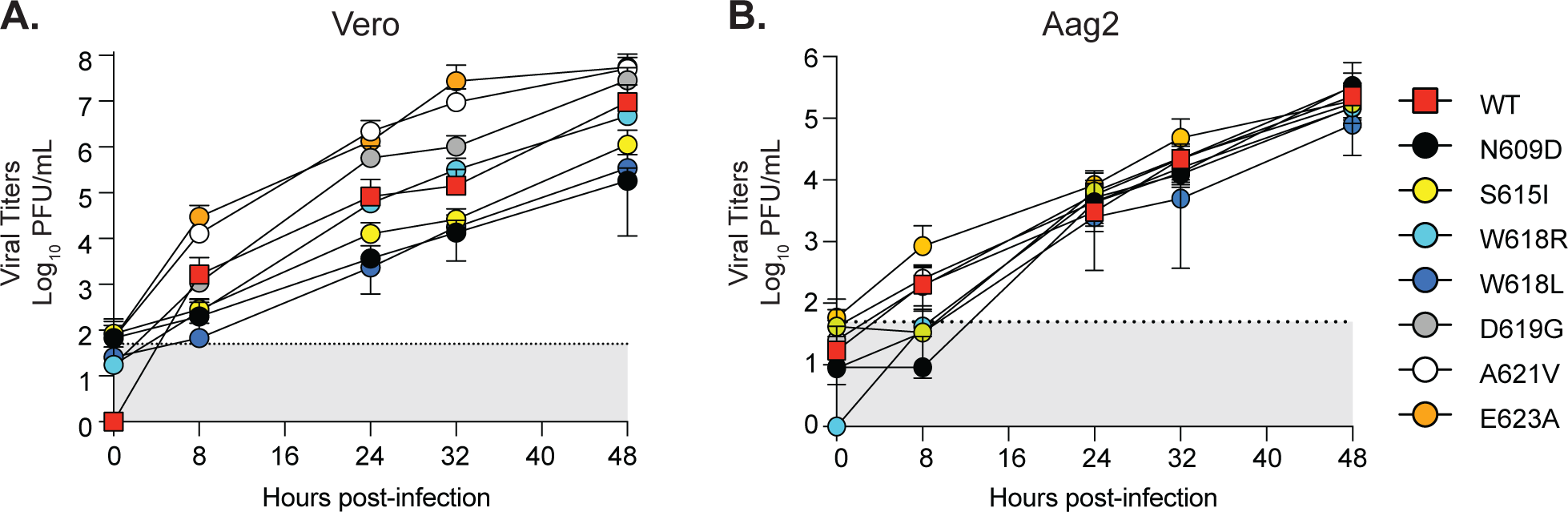
LACV Gc head domain growth in Vero and Aag2 cells. Vero (**A**) and Aag2 (**B**) cells were incubated with each virus at an MOI of 0.1 for 1 hour at 37°C. After incubation, cells were washed, and complete media was added to each well. The supernatant was collected at the indicated time point and infectious virus was quantified by plaque assay on Vero cells. Data represent the mean and SD. Three independent experiments, N=6 for each virus.

### LACV Gc head domain is important for replication and infectivity in human cells

We next tested whether the LACV Gc head domain residues were critical for replication in human neurons and myoblasts, relevant cell lines for LACV infection (28, 29) (**Fig. 3**). We infected human neuroblastoma SH-SY5Y cells and human myoblasts with WT LACV or each Gc head domain variant and quantified infectious virus production at 24 hours post-infection. In SH-SY5Y cells, we found that the variants Gc N609D, S615I, and W618L were attenuated in virus growth up to 100-fold compared to WT LACV, similar to the growth in Vero cells (**Fig. 2A**). In human myoblasts, we observed that the attenuated Gc N609D, S615I, and W618L variants in neurons and Vero cells were not as severely impacted. On the other hand, we found that the Gc variants W618R, D619G, A621V, and E623A were able to enhance infection, similar to what we saw in Vero cells. These results demonstrate that the Gc head domain can influence virus replication in human cells.

**Figure 3:**
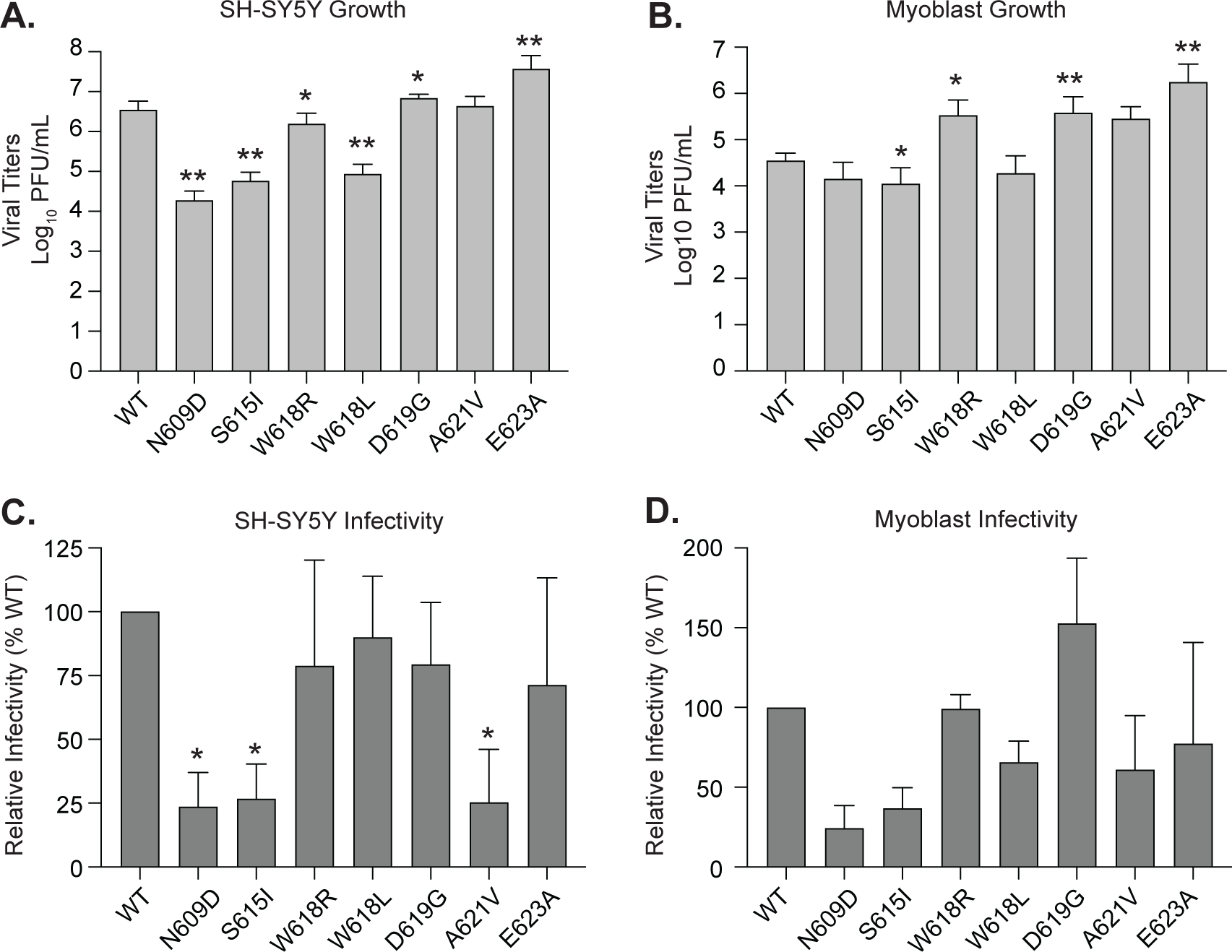
LACV Gc head domain variant infectivity and growth are attenuated in human cells. **(A and B)** Human SH-SY5Y (**A)** or myoblasts (**B**) were incubated with each virus at an MOI of 0.1 for 1 hour at 37°C. After incubation, cells were washed, and complete media was added to each well. The supernatant was collected at 24 hours post-infection and infectious virus was quantified by plaque assay on Vero cells. Data represent the mean and SD. Three independent experiments. N=6/virus. Mann-Whitney test. *p < 0.05, **p< 0.01. (**C and D**) Human SH-SY5Y (**C**) or myoblasts (**D**) were incubated with each virus at an MOI of 1 for 1 hour at 37°C. Following incubation, complete media containing 20 mM NH_4_Cl was added and the cells incubated for 24 hours at 37°C. Cells were then fixed, stained with LACV anti-sera and DAPI, and the number of infected cells were quantified by microscopy. Data are normalized to WT LACV infection. Data represent the mean and SD of three (myoblasts) or four (SH-SY5Y) independent experiments (N=3-4). Mann-Whitney test *p< 0.05.

Given our plaque size phenotypes (**Fig. 1C**) and the location of each residue on the Gc spike, we hypothesized these residues would be important for virus infectivity. To test this hypothesis, we performed infectivity assays by incubating WT LACV and each variant with human neurons or myoblasts for one hour to allow entry. We then added media containing 20 mM ammonium chloride to stop future virus spread, allowing us to only address the initial infection (**Fig. 3C and D**). In both neurons and myoblasts, we found that the Gc variant N609D and S615I had reduced virus infection indicating that these variants may have defects in virus entry. Interestingly, the Gc variant A621V was also attenuated in the infectivity assay (**Fig. 3C**) yet showed WT levels of infectious particle production in growth assays (**Fig. 3A**). These results may suggest that the Gc A621V variant could have defective cell entry, rescued by advantages in virus replication, assembly, or egress. Taken together, we conclude that the LACV Gc head domain is critical for virus entry and replication. Moreover, specific residues within the head domain can significantly attenuate or enhance virus replication.

### The Gc head domain variant N609D attenuates LACV virulence and dissemination in mice

The Gc head domain variant N609D displayed the strongest phenotypes *in vitro* with reduced replication and infectivity. These results allowed us to hypothesize that the Gc N609D variant may significantly reduce virulence. To test this hypothesis, we infected three-week-old wild-type C57BL/6J mice with WT LACV or the Gc N609D via the footpad and weighed mice daily for two weeks (**Fig. 4A and B**). We found that while all of the mice infected with WT LACV succumbed to infection by eight days, the mice infected with Gc N609D all survived the infection (**Fig. 4A**). Importantly, the mice infected with Gc N609D generated antibodies that neutralized both WT LACV and the Gc N609D variant (**Fig. 4C and D**), indicating that they were infected.

**Figure 4:**
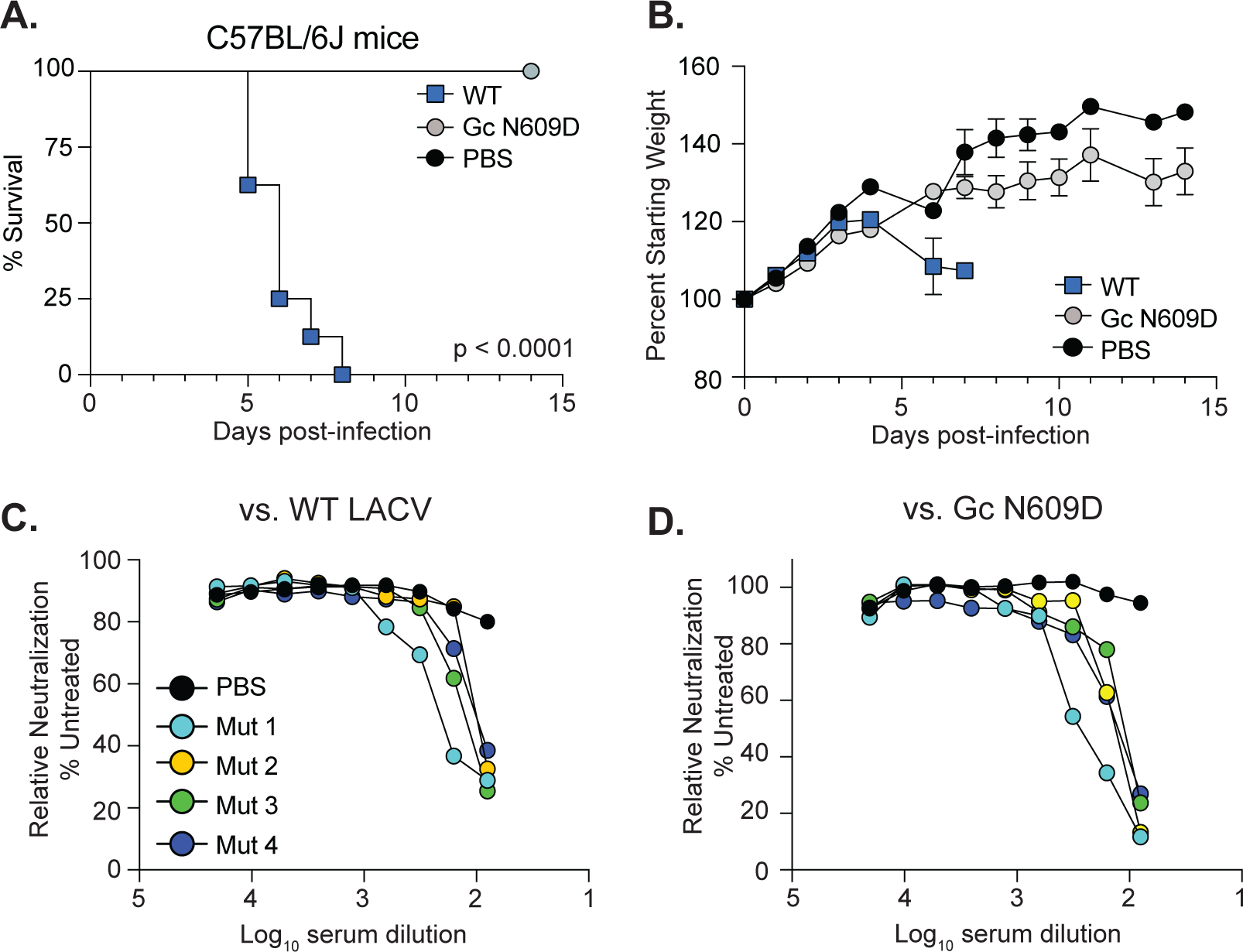
The Gc variant N609D is attenuated in wild-type mice. **(A and B)** Three-week-old male and female C57BL/6J mice were infected with 20,000 PFU of each virus or PBS via the footpad. Mice were monitored (**A**) and weighed daily (**B**). Data represent two independent infections. N=8 mice for each virus. Mantel-Cox test. (**C and D**). At 14 days post-infection, surviving mice were euthanized and the serum collected to address neutralizing antibodies. Serum was inactivated and 2-fold dilutions were mixed with 5000 PFU of WT LACV (**C**) or the Gc N609D virus (**D**) for 1 hour at 37°C. Following incubation, the virus-serum mix was added to Vero cells for 48 hours. Cells were fixed, stained, and the number of infected cells quantified by microscopy.

Given the complete attenuation of the LACV Gc N609D variant in WT mice, we wondered whether this variant would also be attenuated in a more susceptible mouse model. We infected *Ifnar1^-/-^* mice, which lack the type I interferon alpha receptor, with WT LACV or the Gc N609D variant via the footpad (**Fig. 5**). We observed that all mice infected with WT LACV succumbed to infection at five days post-infection. However, we found that for mice infected with Gc N609D, survival was extended several days, with two out of seven mice surviving the infection (**Fig. 5A and B**). The mice surviving the infection lost weight yet recovered around day seven, and generated potent neutralizing antibodies to WT LACV and the Gc N609D variant (**Fig. 5C and D**).

**Figure 5:**
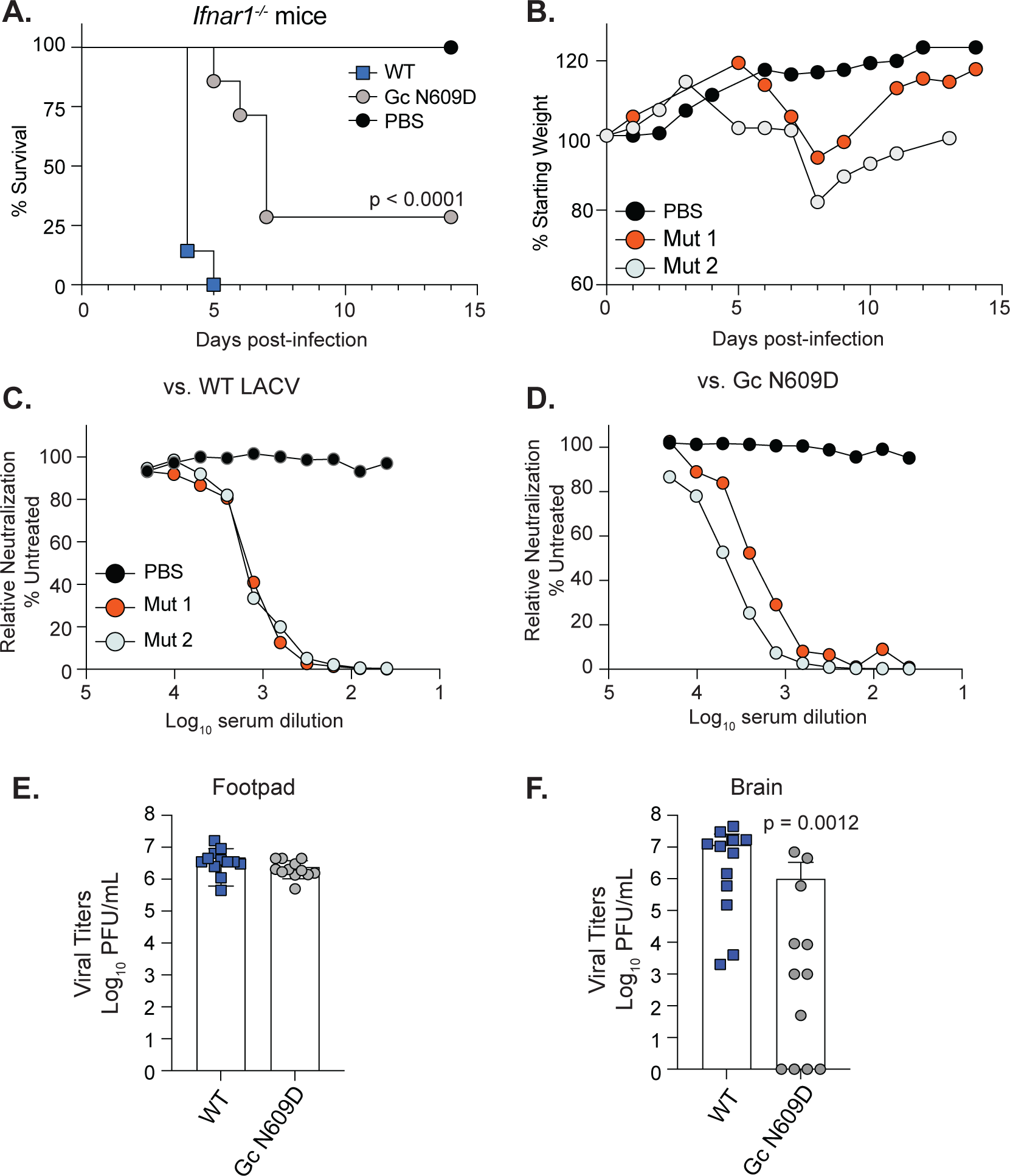
The Gc head domain is important for virulence and dissemination in *Ifnar1^-/-^*. **(A and B)** Three-week-old male and female *Ifnar1^-/-^* mice were infected with 50,000 PFU of each virus or PBS via the footpad. Mice were monitored (**A**) and weighed daily (**B**). Data represent two independent infections. N=7 mice for each virus. Mantel-Cox test. (**C and D**). At 14 days post-infection, surviving mice were euthanized and the serum collected to address neutralizing antibodies. Serum was inactivated and 2-fold dilutions were mixed with 5000 PFU of WT LACV (**C**) or the Gc N609D virus (**D**) for 1 hour at 37°C. Following incubation, the virus-serum mix was added to Vero cells for 48 hours. Cells were fixed, stained, and the number of infected cells quantified by microscopy. (**E and F**). Three-week-old male and female *Ifnar1^-/-^* mice were infected with 20,000 PFU of each virus or PBS via the footpad. At three days post-infection, mice were euthanized, and the footpad (**E**) and brain (**F**) were collected, homogenized, and infectious virus was quantified by plaque assay. Data represent the mean and SD of two independent experiments. N=12 mice for each virus. Mann-Whitney test. p value is shown.

Finally, we hypothesized that the attenuation of the GC N609D variant in *Ifnar1^-/-^* mice was due to not being able to disseminate from the site of infection to the brain. To test this hypothesis, we infected *Ifnar1^-/-^* mice with WT LACV and the Gc N609D virus and addressed viral titers in the footpad (site of infection) and brain at three days post-infection. In the footpad (**Fig. 5E**), both viruses replicated to similar levels indicating that the Gc N609D variant is capable of infecting cells and replicating at the sight of infection. However, when we looked at viral titers in the brain at three days post-infection, we found that while WT LACV replicated to high titers, many of the mice infected with the Gc N609D variant had significantly reduced viral titers (**Fig. 5F**) with several mice having no detectable infectious virus in the brain. These results indicate that Gc head domain, specifically residue N609, is important for virus dissemination and pathogenesis.

### The Gc head domain conservation across LACV lineages and other orthobunyaviruses

Our *in vivo* evolution studies identified the Gc head domain as a potential hotspot for adaptation (15). Therefore, we asked whether the LACV Gc head domain has been changing over time and between LACV lineages. We aligned the LACV Gc head domain regions (amino acids 477 to 722) of all complete LACV M segment sequences (31 in total) deposited in the NCBI database (**Supplemental Figure 1**). For LACV Lineage I, we found that the Gc head domain remained mostly constant with only two mutations present in over 60% of deposited sequences (V528I and K548R; **Table 1**, **Fig. 6A**). These two residues are found at the base of the head domain away from the mutations we had found in our *in vivo* study. In LACV Lineage II, there were six mutations with five localized to the tip of the head domain (**Table 2**, **Fig. 6A**). Finally, in LACV Lineage III, which is the most divergent lineage (30), there were 19 amino acid changes that largely localized along the side of the head domain and at the base (**Fig. 6A**).

**Figure 6:**
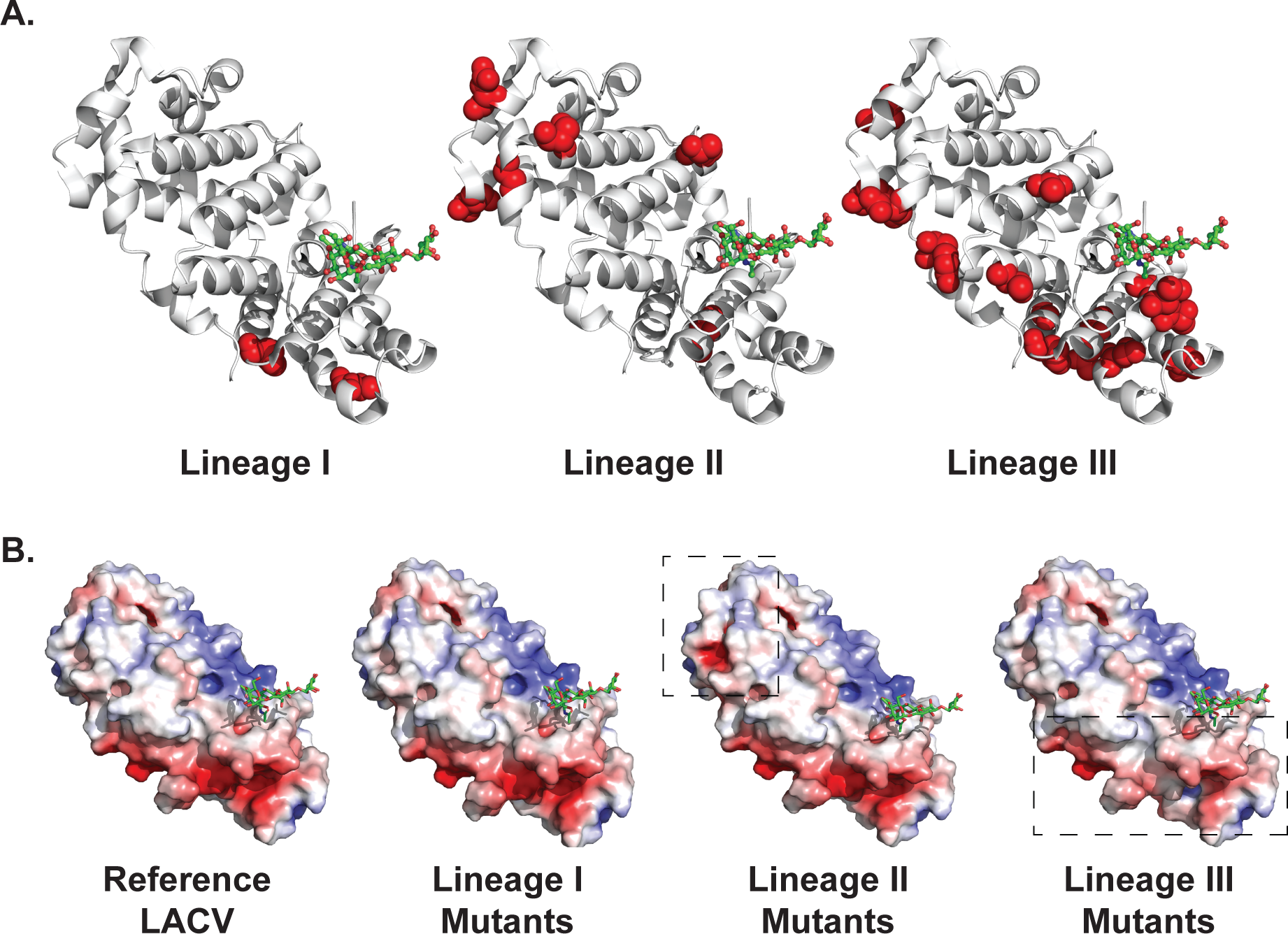
Changes in the Gc head domain across LACV lineages. **A.** Natural mutations found in the Lineage I, II, and III Gc head domains are shown in red. The prototypical LACV/Human/MN/1960 strain was used as a reference and mutations were considered major changes if they were found in >60% of the sequences available. **B.** Electrostatic potentials of the reference LACV head domain (PBD: 6H3W) and the Lineage I, II, and III natural Gc head domain variants. Negative electrostatic potential is in red and positive electrostatic potential is in blue.

**Table 1:**
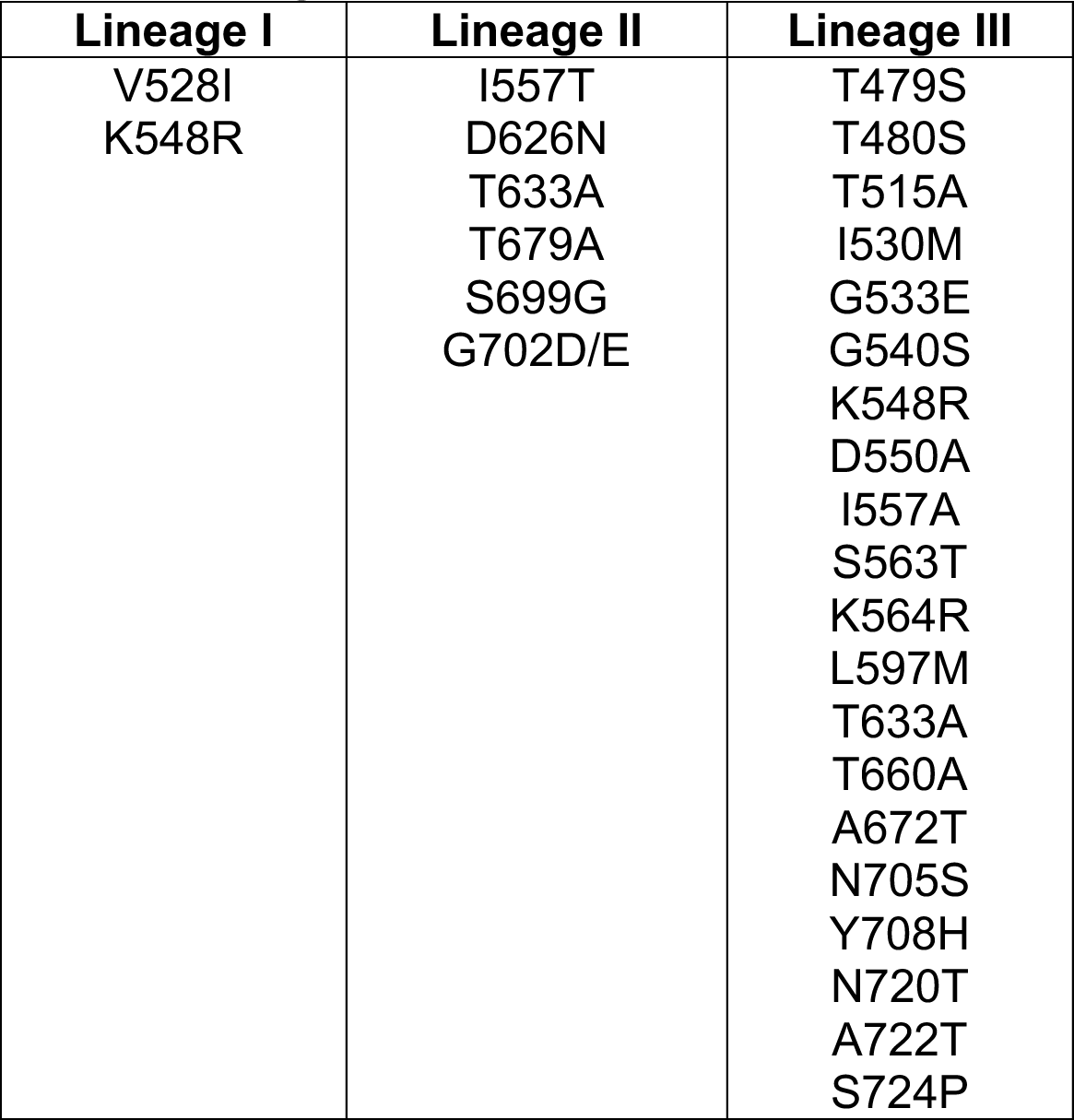
LACV Gc head domain mutations across lineages.

**Table 2:**
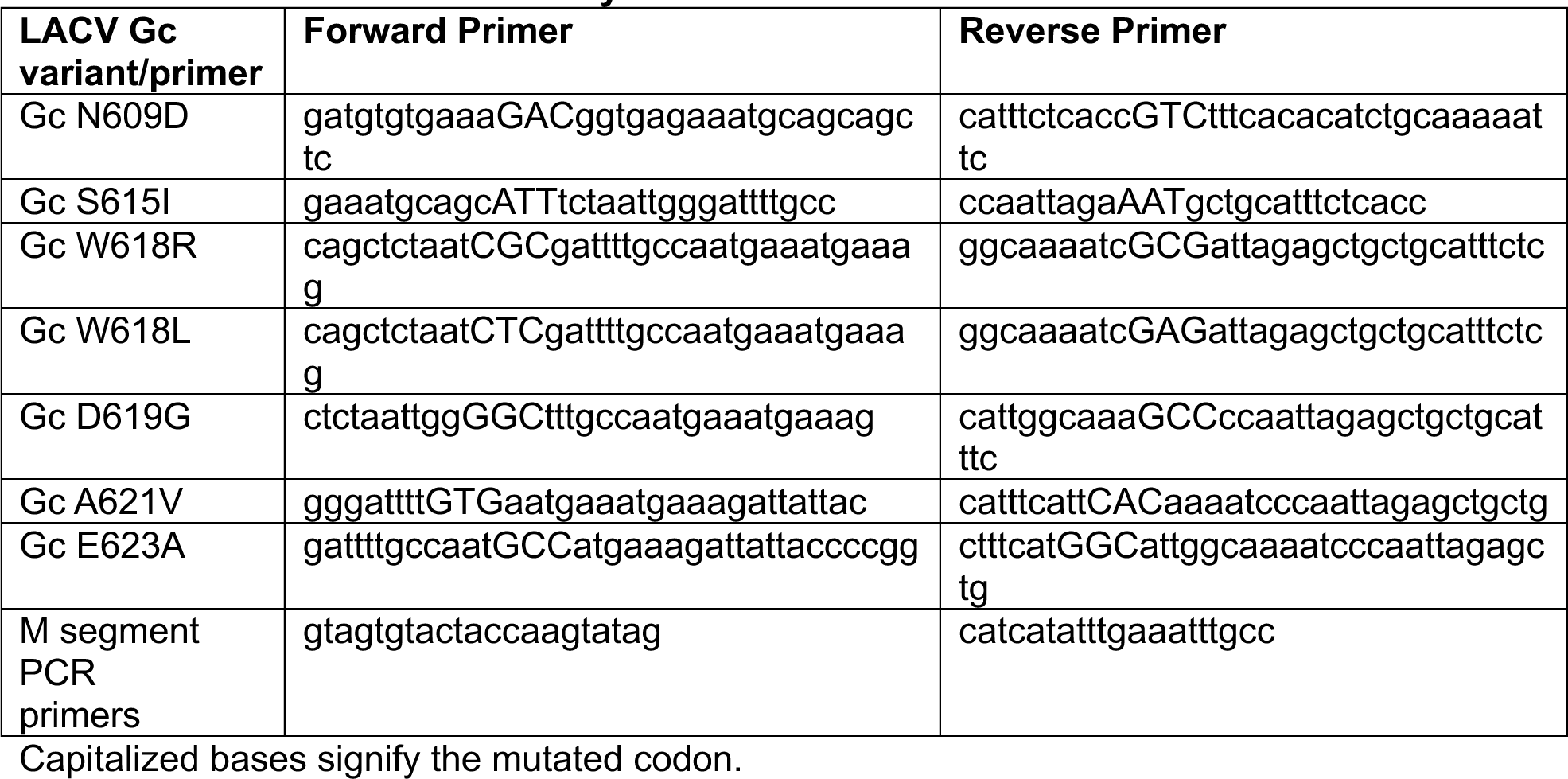
Primers used in this study.

Looking closer at the residues between lineages, many of the changes in the Gc head domain corresponded to changes in charge leading us to hypothesize that these may alter the overall charge of the Gc head domain. To address this hypothesis, we first looked at the overall charged distribution of the LACV Gc head domain (**Fig. 6B**). The Gc head domain has a largely negative charged region running along the side of the head domain with a negative charged spot on top and a positive charged region on the side (**Fig. 6B**). When we introduced changes for lineage I, II, and III into the LACV head domain structure we found that these variants changed the overall charge of the Gc head domain both at the tip, the side, and the base of the head domain (**Fig. 6E, boxes**). These results suggest that alterations in the Gc head domain can influence charged patches on the protein surface that are critical for protein-protein interactions within the virus or host-pathogen interactions needed for infection.

Finally, we asked whether the Gc head domain residues we studied in LACV were conserved across other orthobunyaviruses (**Fig. 7**). Looking at the head domain of the related orthobunyaviruses OROV and BUNV, we found that there are structurally similar alpha helices making up the top portion of the head domains (**Fig. 7A, colored helices**). Moreover, there are similar amino acid residues present in the top of the orthobunyavirus head domains including a conserved glutamine which we showed in LACV is critical for pathogenesis as well as a conserved aromatic tryptophan or tyrosine. Interestingly, both LACV and OROV contain an identical glutamic acid at position 623 which we have shown is capable of enhancing LACV infection *in vitro*.

**Figure 7:**
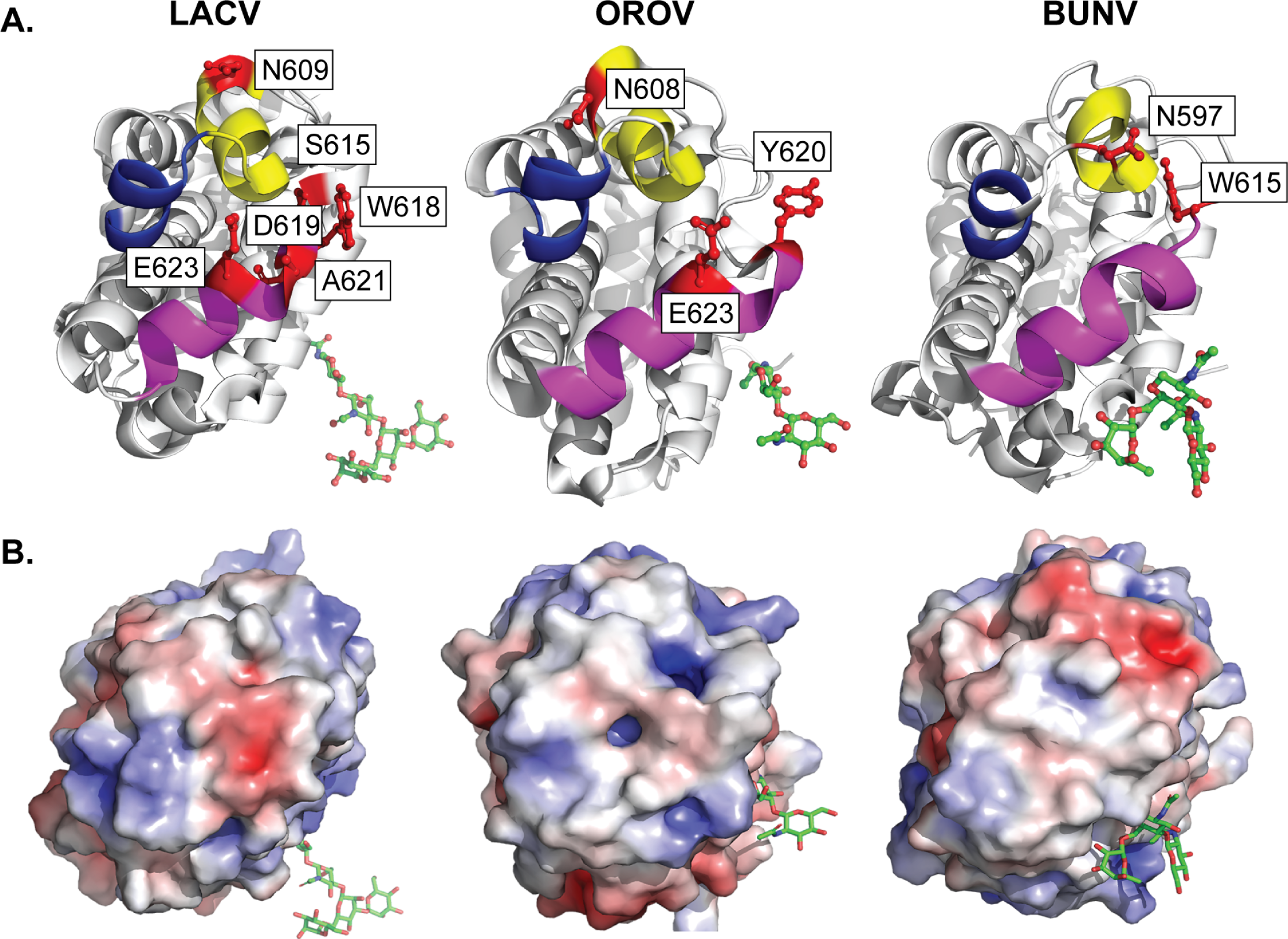
Conservation of the Gc head domain among orthobunyaviruses. **A.** Crystal structure of the LACV (PDB: 6H3W), OROV (PDB: 6H3X), and BUNV (PDB: 6H3V) Gc head domains with residues shown in red. **B.** Electrostatic potential of the LACV, OROV, and BUNV Gc head domain tip.

While there are similarities between orthobunyavirus Gc head domains at the amino acid level, we found major differences in the charge distribution of the Gc head domain between viruses (**Fig. 7B**). These results suggest that the location of charged residues could impact critical host-pathogen interactions necessary for individual virus biology.

## Discussion

Orthobunyaviruses are significant human pathogens capable of devastating outbreaks. These viruses encode Gc, a class II fusion glycoprotein, that is critical for virion assembly and entry. The Gc glycoprotein is functionally and structurally similar to those of alpha- and flaviviruses in domain II which includes the fusion loop. However, orthobunyaviruses are unique in that they encode a variable amino-terminal head domain that forms the tip of the virion trimeric spike. Our understanding of how the head domain functions is not well defined.

In this study, we took advantage of *in vivo* evolution of La Crosse virus (LACV) in mice and mosquitoes where we identified several mutations clustering in the Gc head domain. Given the location of these changes and that several of them are found in nature, we hypothesized these are important for LACV and orthobunyavirus biology. We generated each variant and tested how these mutations influenced virus growth in multiple cell lines. We found that while these variants were largely genetically stable in mammalian cells, they reverted to the wild-type residue in mosquito cells suggesting a strong selective pressure in these cells. These results are interesting as we identified several of these variants in mosquitoes. One explanation for these results could be that our previous *in vivo* evolution studies used the LACV Lineage I strain Human/MN/1960 which differs from our infectious clone Lineage I system Mosq/NC/1978 by 29 amino acids in the M segment. Regardless, we find that specific head domain residues could both influence replication in human neurons and myoblasts, indicating an important role for the head domain virus replication and spread. Interestingly, replication may be cell type specific as while we find several of the variants attenuated in neurons, we do not see the same results in myoblasts. Moreover, although we find that the variants W618R, D619G, A621V, and E623A can change replication in myoblasts, this does not seem to be due to changes in infectivity. These results may suggest that these variants are important for other steps in the viral life cycle to be explored in the future. Our results show that even single amino acid changes can have major impacts on replication highlighting the importance of the Gc head domain in orthobunyavirus biology.

In addition to roles *in vitro*, we found that the head domain and specifically residue N609 were important for virulence and dissemination in mice. In WT mice, the Gc head domain variant N609D was completely attenuated, yet the mice started to lose weight around seven days post-infection and retained this weight for the remainder of the experiment. These results along with neutralizing antibodies show that the mice are infected, yet do not succumb to infection. Moreover, we find that in the highly susceptible type-I interferon deficient mice, the Gc N609D variant is attenuated although mice do succumb to infection. In the two *Ifnar1^-/-^* mice that survived the infection, both lost weight, recovered, and produced neutralizing antibodies which we hypothesize are critical for viral clearance. Looking at virus dissemination, we find that while the Gc N609D variant can replicate to WT levels at the site of infection, it has defects in neuroinvasion from the site of infection to the brain. Similar defects in neuroinvasion have been seen for LACV fusion loop mutants (31), however these variants also showed defects in the muscle suggesting a global defect in virus entry. Our in vivo and in vitro results suggest that the Gc head domain is critical for cell-specific interactions and entry mechanisms that may drive virus dissemination. Together, these results show that the Gc head domain plays an essential role in LACV virulence through dissemination and suggest that if the virus can be restricted to the periphery for long enough that antibodies are produced, the host can recover.

Finally, we see that the LACV Gc head domain, while variable between lineages is largely conserved at key residues at the tip of the spike. Given their location, we hypothesize that these residues may be interacting with an unknown LACV receptor or other host factors on the cell surface to facilitate entry. In the case of the Gc N609D variant, we speculate that these interactions are disrupted, leading to reduced infection or infection via an alternative pathway which attenuates virus dissemination and overall pathogenesis. An additional hypothesis may be that these Gc head domain residues are important for proper spike assembly and therefore changes in glycoprotein structure can change host-pathogen interactions needed for virus entry. Specifically, the presence of distinct positive charged patches may suggest potential interactions with negatively charged glycosaminoglycans as is the case for other arboviruses (32–34). Changes in the charge network of the Gc head domain could explain why the different lineages of LACV differ in pathogenesis (35). Future work will be important to investigate the role of the Gc head domain in the pathogenesis of LACV and other orthobunyaviruses. We find that the OROV Gc head domain for example is structurally similar to that of LACV with similar residues maintained at the Gc tip. It will be important to interrogate how the head domain of orthobunyaviruses facilitates virus dissemination and disease to better understand how these pathogens establish infections.

## Acknowledgements

We thank all members of the Stapleford Lab for helpful discussions on this manuscript. This work was supported by funding from the New York University Grossman School of Medicine Startup and NIAID/NIH R01 AI162774-01A1.

## Materials and Methods

### Cells

Vero cells (CCL-81, ATCC) were grown in Dulbecco’s Modified Eagle Media (DMEM) supplemented with 10% newborn calf serum (NBCS). BHK-21 BSR/T7 cells (36), a gift from Dr. Steven Whitehead at the National Institutes of Health (NIH) were grown in DMEM supplemented with 10% fetal bovine serum (FBS), 1% non-essential amino acids (NEAA), 10 mM HEPES and 1 mg/mL Geneticin added every other passage to maintain T7 selection. Human neuroblastoma cells (SH-SY5Y, CRL-2266, ATCC) were grown in a 50:50 mix of Eagle’s Minimum Essential Media (ATCC) and F12 Medium supplemented with 10% FBS. Immortalized human myoblasts were a gift from Dr. Michael Kyba at the University of Minnesota(28). Myoblasts were grown in HAM’s/F10 Nutrient Mixture supplemented with 20% FBS, 1x Glutamax, 10 ng/mL human basic fibroblast growth factor, 40 ng/ml dexamethasone, and 100 μM beta-mercaptoethanol. All mammalian cells were maintained at 37°C with 5% CO_2_. *Aedes aegypti* cells (Aag2) were a gift from Dr. Paul Turner at Yale University and maintained in DMEM supplemented with 10% FBS, 1% NEAA, and 10 mM HEPES at 28°C with 5% CO_2_. All cells were confirmed mycoplasma free by monthly testing.

### Viruses

The La Crosse virus (LACV) infectious clone system was obtained from Dr. Whitehead (27). Gc head domain mutants were generated by site-directed mutagenesis of the M segment using the primers in **Table 2**. All plasmids were sequenced at Plasmidsaurus to ensure there were no second-site mutations. To generate each virus, BHK-21 BSR/T7 cells were transfected with 2 μg of each of the S, M, and L plasmids using Trans-IT LT1 transfection reagent (Mirus). 24 hours post transfection, media was replaced, and cells incubated at 37°C for five days. Supernatants were collected, aliquoted, and stored at - 80°C. To generate working virus stocks, virus from each transfection was amplified on a monolayer of Vero cells. Viruses were collected, centrifuged at 1,200 rpm for 5 minutes, aliquoted, and stored at −80°C. Viral titers were quantified by plaque assay as described below.

### RNA extractions and Sanger sequencing

RNA was extracted using Trizol (Thermo) and the Direct-Zol plus RNA extraction kit (Zymo Research) following the manufacturer’s instructions. cDNA was generated using the Maxima H Minus first-strand cDNA synthesis kit (Thermo) and used for Phusion PCR (Thermo) with the M segment sequencing primers in **Table 2**. PCR amplicons were purified with the Macherey-Nagel PCR clean up kit and Sanger sequenced at Plasmidsaurus. PCR sequences were aligned to the infectious clone reference using SnapGene (version 8.0.3).

### Plaque assay

350,000 Vero cells were seeded in 12-well plates and incubated with 10-fold dilutions of each virus for 1 hour at 37°C. 0.8% agarose in DMEM containing 2% NBCS and 1x antibiotic/antimycotic (Gibco) was added and cells incubated for 72 hours. Following incubation, cells were fixed with 4% formalin for 1 hour, agarose plugs removed, and cells stained with crystal violet. Viral titers were quantified by counting the number of plaques on the lowest countable dilution. Plaque size was quantified using Image Lab (BioRad; version 6.1.0).

### LACV growth curves

Vero cells (55,000 cells/well), Aag2 cells (200,000 cells/well), myoblasts (55,000 cells/well), and SH-SY5Y cells (200,000 cells/well) were seeded in Poly-L-Lysine coated 24-well plates. Cells were incubated with each virus at a MOI of 0.1 for 1 hour at 37°C or 28°C. Virus was removed, cells were washed twice with phosphate buffered saline (PBS), and complete media added. Supernatant was collected at the indicated time points and infectious viral titers were quantified by plaque assay as described above.

### LACV infectivity assays and immunostaining

Myoblasts (10,000 cells/well) and SH-SY5Y cells (50,000 cells/well) were seeded in black 96-well Costar clear bottom plates. Cells were washed once with PBS and incubated with each virus at an MOI of 1 for 1 hour at 37°C. After incubation, media containing 20 mM ammonium chloride was added to block virus spread. Cells were incubated for 24 hours and fixed in 4% paraformaldehyde (PFA). For LACV staining, cells were then washed three times with perm/wash, incubated with 0.25% Triton for 10 minutes, and incubated in blocking buffer (0.2% bovine serum albumin, 0.05% saponin in PBS) for 1 hour at room temperature. Cells were then incubated with a 1:2000 dilution of primary rabbit anti-LACV antibodies (a gift from Dr. Karin Peterson at the NIH) in blocking buffer for 2 hours, washed extensively with perm/wash, and incubated with a 1:10,000 dilution of secondary goat anti-rabbit IgG Alexa488 and DAPI (1:1000 dilution) for 1 hr. Cells were washed three times with perm/wash and PBS was added. The number of infected cells was quantified on the CX7 Cell Insight high content microscope.

### Mouse infections

All animal work was completed at the NYU Grossman School of Medicine under IACUC protocol IA16-01783. For survival assays, three-week-old wild-type C57BL/6J (Strain # 000664; Jackson Laboratory) and *Ifnar1^-/-^* mice (Strain #028288; Jackson Laboratory) were infected with 20,000 or 50,000 PFU of each virus in the left rear footpad while under anesthesia with isoflurane. Mice were weighed daily and euthanized when the weight reached less than 20% of the starting weight. All mice were euthanized after 14 days. For viral titers, mice were infected with 20,000 PFU and euthanized at 3 days post-infection. The footpad and brain were harvested in 750 μL of plaquing media containing steel beads. The tissue was homogenized for 5 minutes at the setting 12 using a Bullet Blender Storm Pro (Next Advance) and clarified by centrifugation. Viral titers were quantified by plaque assay as described above.

### LACV neutralization assays

At 14 days post-infection, mice were euthanized and serum was collected. Serum was inactivated at 56°C for 30 minutes and 2-fold dilutions made in DMEM containing 2% NCBS. Each dilution was mixed with 5000 PFU of WT LACV or the Gc N609D virus and incubated at 37°C for 1 hour. Following incubation, the virus mix was added to Vero cells (15,000 cells/well in a black 96-well Costar clear bottom plate) and incubated for 48 hours at 37°C. Cells were then fixed with 4% PFA, stained for LACV antigen, and the number of infected cells were quantified with the CX7 CellInsight High Content microscope as described above.

### Protein structures and alignments

PyMOL (Version 3.1.1) was used for the structures of the head domains of LACV (PDB: 6H3W), OROV (PDB: 6H3X), and BUNV (PDB: 6H3V). The PyMOL mutagenesis wizard was used to introduce mutations and APBS Electrostatics plugin was used for analyzing charge distribution. SnapGene (Version 8.0.3) was used to align all LACV M segment complete protein sequences found in NCBI.

### Statistics

All data were analyzed using GraphPad Prism (Version 10.4.2). Data are represented as the mean and standard deviation. *In vitro* experiments were completed three independent times with internal duplicates where noted. *In vivo* experiments were completed at least two independent times with the number of animals found in the figure legends. P < 0.05 is considered statistically significant.

**Supplemental Figure 1:** Alignment of LACV M segment amino acid region 477-725. Lineages are depicted to the left of the accession numbers. White shading signifies amino acid change from reference LACV/Human/MN/1960.

